# Cycling hypoxia selects for constitutive HIF stabilization

**DOI:** 10.1101/2020.10.28.359018

**Authors:** Mariyah Pressley, Jill A. Gallaher, Joel S. Brown, Michal R. Tomaszewski, Punit Borad, Mehdi Damaghi, Robert J. Gillies, Christopher J. Whelan

## Abstract

Tumors experience temporal and spatial fluctuations in oxygenation. Hypoxia inducible transcription factors (HIF-α) in tumor cells are stabilized in response to low levels of oxygen and induce angiogenesis to re-supply oxygen. HIF-α stabilization is typically facultative, induced by hypoxia and reduced by normoxia. In some cancers, however, HIF-α stabilization becomes constitutive even under normoxia, a condition known as *pseudohypoxia*. Herein, we develop a mathematical model that predicts the effects of fluctuating levels of oxygen availability on stabilization of HIF-α and its client proteins based on fitness. The model shows that facultative regulation of HIF-α always promotes greater cell fitness than constitutive regulation. However, cell fitness is nearly identical regardless of HIF-α regulation strategy when there are rapid periodic fluctuations in oxygenation. Furthermore, the model predicts that stochastic changes in oxygenation favor facultative HIF-α regulation. We conclude that rapid and regular cycling of oxygenation levels selects for pseudohypoxia.

## Introduction

Phenotypic plasticity, the production of alternative phenotypes in response to variable environments, is ubiquitous in nature^1^. Phenotypic plasticity confers flexibility that allows an organism to survive in the face of often unpredictable and rapid changes in its environment. Many organisms, from microbes to humans, vary gene expression facultatively. In this way phenotypic expression matches demand^2,3^. In contrast, when environments are constant, or are predictably variable, an intermediate and constitutive level of expression may be favored over facultative expression^4,5^.

Inducible or facultative defenses, such as defensive chemicals in plants, and spines and projections in zooplankton such as rotifers and cladocerans, include remarkably diverse and well-studied examples of phenotypically plastic responses to biotic and abiotic threats or stressors. Biotic threats include herbivores, predators, pathogens and parasites. An important abiotic threat is hypoxia, a reduction in oxygen availability. Hypoxia may be either acute, intermittent, or chronic, and it may be experienced at the organismal, tissue, or cellular levels. Cells respond to hypoxia via stabilization of the Hypoxia-inducible Factor (HIF), an inducible defense against both acute and chronic hypoxia within the cellular environment. HIF is a heterodimeric transcription factor that induces expression of genes that lead to tissue re-oxygenation. The evolution of inducible defenses, like HIF, appears to be favored by unpredictability of environmental conditions, reliable cues of those conditions, and a high cost of the defense^6-8^.

Oxygen levels in normoxic or hypoxic tissues encompass a wide range of values depending on several factors, including gender, time of day, tissue type, and degree of vascularization^9,10^. In tumors, significant heterogeneity in oxygen levels result from both a dynamic ecosystem of blood vessels of varying functionality, and cancer cells with different tolerances to hypoxia. Within a nascent tumor ecosystem, cancer cells, which can somatically evolve, experience both acute and chronic hypoxia due to rapid growth, limited blood supply, and disorganized vascular delivery systems^11^. This leads to complex cycles of oxygenation and hypoxia, characterized as “waves” and “tides”^12-14^. Thus, an intermittent or temporal instability in oxygen supply is a cardinal feature of tumors. We have proposed that this generates strong evolutionary selection pressures for more aggressive cancer cell phenotypes^15^.

Like nearly all metazoan cells, cancer cells possess mechanisms to respond to heterogeneity in the supply of oxygen, including activation of a family of hypoxia-inducible transcription factors (HIF-1α, HIF-2α, and HIF-3α; hereafter HIF-α), cell cycle arrest, a coordinated decrease in oxidative phosphorylation with an increase in glycolysis (Pasteur Effect), and the secretion of angiogenic factors to promote blood vessel formation^16-21^. HIF-α is ubiquitously and continuously expressed in all cells. In well-oxygenated environments (normoxia), HIF-α is hydroxylated and ubiquitinated and is thus recognized and degraded by proteasomes^19^. In an oxygen-depleted state, hydroxylation and hence, degradation of HIF-α is inhibited, promoting the transcription of genes that regulate proliferation^22,23^, cellular metabolism^19^, angiogenesis^24^ and erythropoiesis^25-26^. If well-regulated, the recruitment of blood vessels to the site of HIF-α stabilization increases the supply of oxygen. Once O_2_ levels return to normal (normoxia), HIF-α returns to baseline levels. It has also been observed that aggressive cancers constitutively express hypoxia-related proteins (HRPs) even in the presence of oxygen, a condition known as *pseudohypoxia*^27^. The most common manifestation of this phenotype is the fermentation of glucose under normoxia, known as “aerobic glycolysis” or the “Warburg Effect”^28-30^.

Maintenance of HIF-α levels under cycling hypoxia involves tradeoffs. Under prolonged hypoxia without stabilization of HIF-α, cells die. In contrast, HIF-α stabilization under normoxia comes at a cost. Accumulation of HIF-α in well-oxygenated environments costs energy and resources for the synthesis of HIF-α client proteins that may not be necessary for survival, including activating energetically inefficient glycolysis and expression of the exofacial acidifying pH-stat, carbonic anhydrase isoform 9, CA-IX^31^. As seen in the development of some tumors, regulation of HIF-α switches from a facultative state, where the environment induces the changes in regulation, to a constitutive state, where HIF-α and/or HIF-α client proteins remain above baseline regardless of the environment^15,27^. As this phenotype is associated with cancer progression and aggressiveness, understanding the microenvironmental conditions that select for pseudohypoxia is fundamentally important.

Here, we develop a mathematical model with the goal of determining a cancer cell’s optimal level of HIF-α expression with respect to differences in fluctuating levels of oxygen availability within tumor microenvironments. Specifically, we seek to determine what tumor conditions may cause the evolution of constitutive HIF-α regulation (“hard wired” HIF-α stabilization) from facultative regulation. To do so we compare the maximal expected payoff (net growth rate) between facultative and constitutive HIF-α regulation in environments with different oxygen profiles. We hypothesize that predictable and rapid cyclic fluctuations from normoxia to hypoxia will favor constitutive HIF-α stabilization based on similar ideas in optimal defense theory^4,32^.

Our model investigates how cells may respond to changes in oxygenation with HIF-α expression. In a perfect world, cells would instantaneously optimize HIF-α levels in response to fluctuating oxygen concentrations. As the environment shifts from normoxia to hypoxia, cells would immediately accumulate HIF-α, and vice-versa. However, attaining the appropriate HIF-α level for the current environment is not immediate. There will be time lags in upregulating or down regulating HIF-α. HIF-α production and proteosomal degradation occur continuously^18,27^. When oxygenation levels decline, HIF-α degradation slows, and production permits HIF-α to increase at a relatively slow rate to counter the hypoxic conditions^18^. Upon re-oxygenation, HIF-α can be rapidly degraded. We model cellular regulation of HIF-α as the concentration of oxygen within the tumor and surrounding microenvironment changes temporally – both with regular periodicity and stochastically. The model was informed by empirical evidence for the rates of HIF-α accumulation and degradation as the cells’ microenvironment shifts between normoxia and hypoxia, and vice-versa^33-35^.

## Results

We compared how different HIF-α regulation strategies influence cell fitness under different oxygenation environments. Here, we define a cell’s fitness by its net proliferation rate, or “payoff”. In this model, we assume that the expected payoff depends on the base proliferation rate, the metabolic cost of expressing HIF-α, and the mortality risk of not expressing HIF-α during hypoxia. We use the following expression for a cancer cell’s payoff at time *t, G*(*t*):

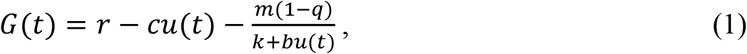

where *r* is the baseline proliferation rate of a cancer cell, *c* is proliferation cost of using HIF-α strategy *u* at time *t, m* is the cell mortality when conditions are hypoxic, *q* is the fraction of time spent in normoxic conditions, *k* is a cell’s intrinsic tolerance to hypoxia in the absence of HIF-α stabilization, and *b* is the benefit of HIF-α expression *u* in reducing mortality when conditions are hypoxic. See Table 1 for definitions, units and values of all parameters used in the models.

**Table 1.**
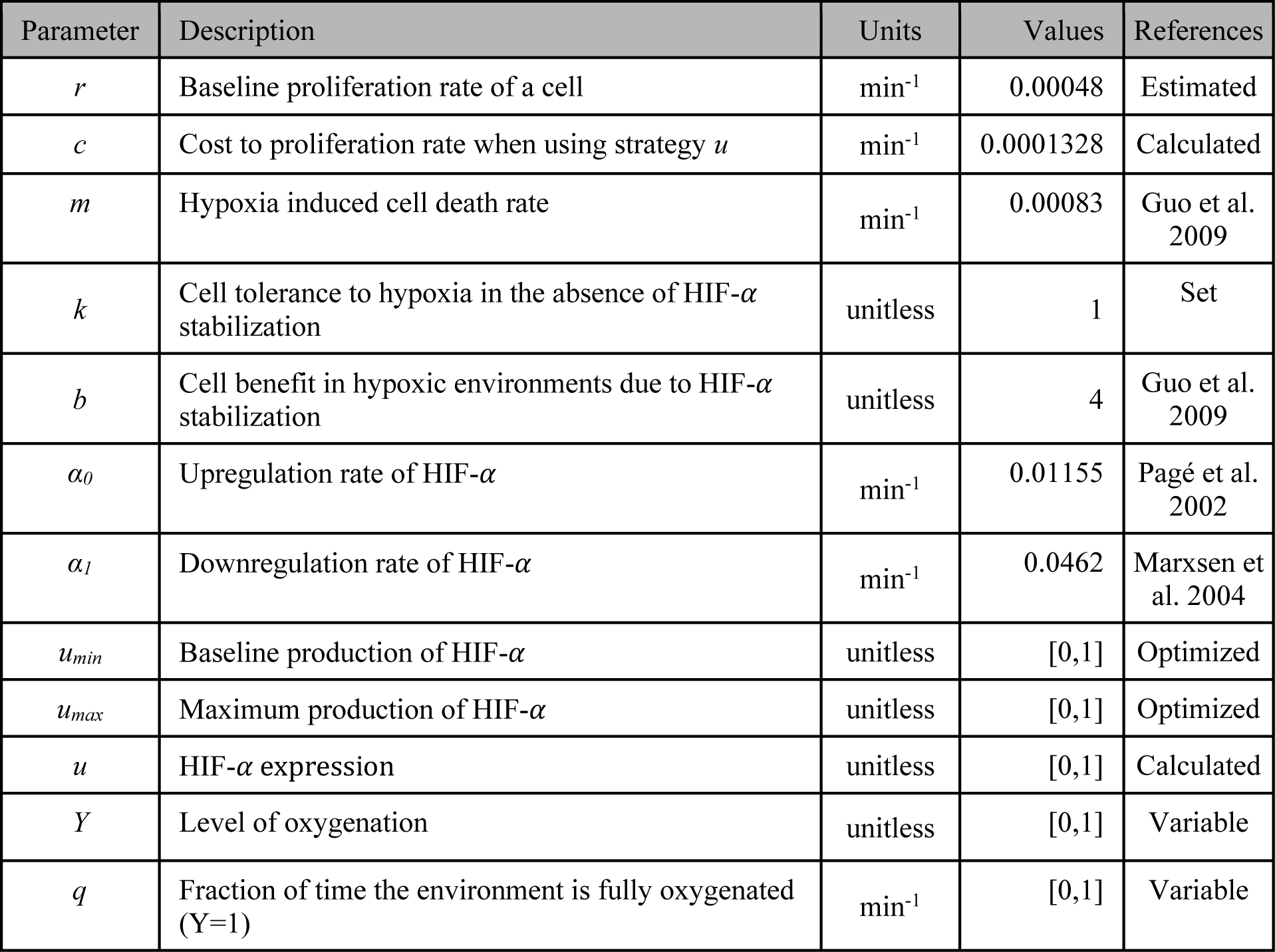
Definitions, units, and values of parameters used in the models. The terms *Y* and *u* are normalized and are therefore unitless. We normalize *k* to 1 because it appears as *m/k* when *u*=0 and appears as *b/k* in *u*\*. The value of *c* is calculated such that *u*\* = 1 when the environment is hypoxic 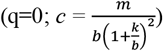.

| Parameter | Description | Units | Values | References |
| --- | --- | --- | --- | --- |
| $r$ | Baseline proliferation rate of a cell | min<sup>-1</sup> | 0.00048 | Estimated |
| $c$ | Cost to proliferation rate when using strategy $u$ | min<sup>-1</sup> | 0.0001328 | Calculated |
| $m$ | Hypoxia induced cell death rate | min<sup>-1</sup> | 0.00083 | Guo et al. 2009 |
| $k$ | Cell tolerance to hypoxia in the absence of HIF-α stabilization | unitless | 1 | Set |
| $b$ | Cell benefit in hypoxic environments due to HIF-α stabilization | unitless | 4 | Guo et al. 2009 |
| $\alpha_0$ | Upregulation rate of HIF-α | min<sup>-1</sup> | 0.01155 | Pagé et al. 2002 |
| $\alpha_1$ | Downregulation rate of HIF-α | min<sup>-1</sup> | 0.0462 | Marxsen et al. 2004 |
| $u_{min}$ | Baseline production of HIF-α | unitless | [0,1] | Optimized |
| $u_{max}$ | Maximum production of HIF-α | unitless | [0,1] | Optimized |
| $u$ | HIF-α expression | unitless | [0,1] | Calculated |
| $Y$ | Level of oxygenation | unitless | [0,1] | Variable |
| $q$ | Fraction of time the environment is fully oxygenated (Y=1) | min<sup>-1</sup> | [0,1] | Variable |

We compare three HIF-α strategies across a variety of oxygenation environments: perfect, constitutive, and facultative. A perfect strategy means instantaneous HIF-α switching in response to normoxic and hypoxic conditions. While this strategy is idealized, it provides a useful point of comparison for the other strategies as a perfect strategy would yield the highest possible payoff. A constitutive strategy assumes that the HIF-α level is constant over time, and a facultative strategy assumes that the HIF-α levels can change at a finite rate in response to oxygen levels. Representative examples of these strategies in a fluctuating environment are shown in the Methods (Fig. 7).

### The perfect strategy

A perfect strategy does not mean perfect fitness in all environments. Mortality and the metabolic costs of HIF-α stabilization means that fitness declines with more time spent in hypoxic conditions. The expected payoff of the perfect strategy (see Eq. (2) in Methods) is shown to decline linearly with the proportion of time spent in hypoxia (Fig. 1A).

**Figure 1.**
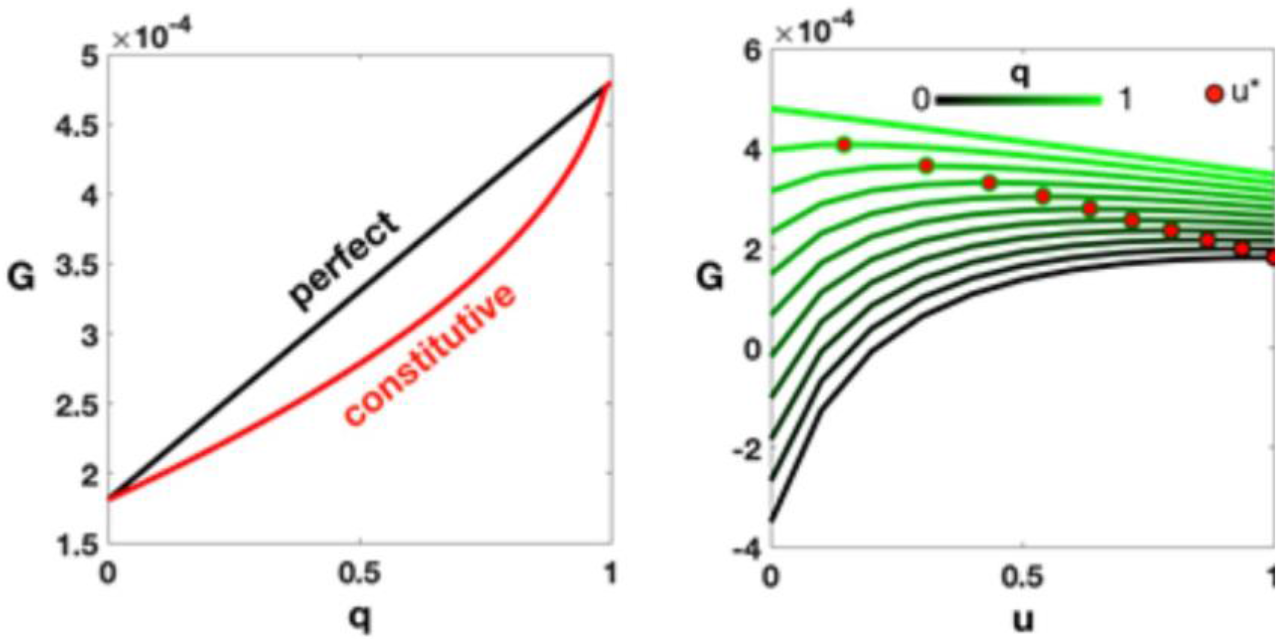
Expected payoffs for perfect and constitutive strategies. A) The payoff (*G*) for perfect and optimal constitutive (*u*\*) strategies for different fractions of time in normoxic conditions, *q*. B) The payoff versus HIF-*α* level, *u*, for different *q*’s. The lines represent all *u* values, while the red dots represent the optimal *u* expression (*u*\*) for each *q* that maximizes the payoff. Parameter values are given in Table 1.

### The constitutive strategy

The optimal constitutive strategy can be found by maximizing the payoff for a constant HIF-α level, *u*\* (see Eq. (3) in Methods). The expression for the payoff can then be analytically solved (see Eq. (4) in Methods). We found that the constitutive strategy payoff is always less than or equal to the perfect strategy payoff (Fig. 1A), being equal only when the environment is always hypoxic (*q* = 0) or always normoxic (*q* = 1). The payoff over different HIF-α regulation strategy values *u* for different fractions of time spent in normoxia *q* is plotted in Fig. 1B, along with the optimal constitutive values *u*^*^. As expected, we find that the payoff is highest when HIF-α expression is lowest (u=0) under constant normoxic conditions. With constant normoxia, as u is increased, the payoff slightly decreases, showing the minor cost of unnecessarily producing HIF-α. However, as conditions become more hypoxic with a low HIF-α expression, the mortality term dominates, and the payoff is drastically reduced into negative values where the cell cannot survive. With constant hypoxia, HIF-α needs to increase in order for *G*≥0 for survival. The optimal values u^*^, shown as red dots, reflect this for all values of *q* in between.

### Facultative expression of HIF-α

With the facultative strategy, *u* changes over time according to Eq. (5) in the Methods. This expression allows *u* to increase in hypoxia and decrease in normoxia at the rates given in Table 1. The rates were based on empirical measurements^33-35^, which found that HIF-α down-regulation occurs about four times faster than HIF-α up-regulation. The optimal facultative strategy involves selecting a baseline *u*_*min*_ and an upper bound *u*_*max*_ that maximizes the expected payoff. For simulations we began with a period of normoxia and a starting value of HIF-α halfway between constitutive expression and the specified *u*_max_. For fixed cycle lengths, the expected payoff converges quickly to a steady state, and we use the payoff in Eq. (6) in the Methods to numerically solve for the optimal lower and upper bounds of *u*\*_*min*_ and an upper bound *u*\*_*max*_.

We compared the optimal facultative response between two different periodicities (either 10 minute or 120 minute intervals) of full oxygenation followed by the same for deoxygenation (*q* = 0.5 in both cases). Initially we compared the payoffs of a constitutive strategy and facultative strategy in the same environment to determine the selection coefficient, which we defined as the fitness advantage for using a facultative strategy (details in the Methods). The selection coefficients over the full range of possible *u*_*min*_ and *u*_*max*_ are shown in Fig. 2A. For longer cycling periods, the facultative strategy has a larger selection coefficient for any given combination of *u*_*min*_ and *u*_*max*_ than for shorter cycle times. Under many circumstances the difference in the selection coefficients for the two strategies are so small as to be negligible. Under such circumstances one might expect the constitutive strategy to prevail.

**Figure 2.**
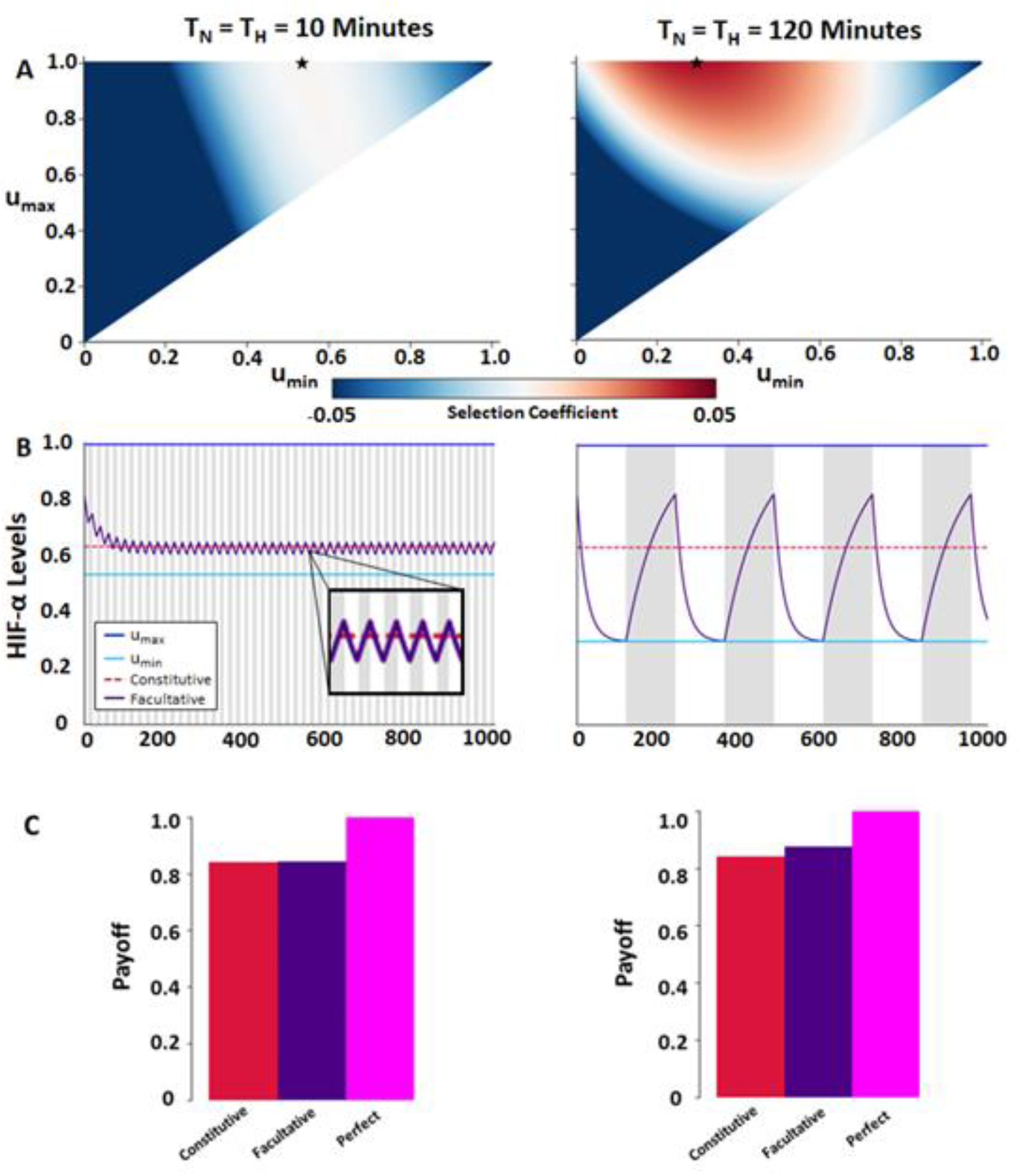
Comparison environments with fixed intervals of short (10 min) and long (120 min) periods of cycling hypoxia. A) Heatmap of the selection coefficients for the facultative strategy for all *u*_*min*_ and *u*_*max*_ combinations. Each star denotes the *u*_*min*_ and *u*_*max*_ combination that maximizes payoffs for the facultative strategy for each fixed interval time. B) HIF-α (u) levels over time using the optimal facultative strategy. The unshaded areas represent periods of full oxygenation while the shaded grey areas represent periods of hypoxia. C) Payoffs for separate strategies of HIF-α expression normalized to the perfect strategy. Parameters are given in Table 1.

We further explored HIF-α expression using the *u*_*min*_ and *u*_*max*_ combination that produced the greatest payoff. When the environment fluctuates with time intervals of 10 minutes, the cell’s payoff was optimized at *u*\*_*min*_=0.540 and *u*\*_*max*_=1. The fluctuation in HIF-α expression occurred rapidly and the *u*\*_*min*_, while always less than, came close to the constitutive value, while *u*\*_*max*_ remained at 1 (Fig. 2B). By increasing time intervals to 120 minutes, the cell’s payoff was maximized at *u*\*_*min*_=0.305 and *u*\*_*max*_=1. The longer cycle times resulted in larger fluctuations in HIF-α expression and a greater superiority of the facultative strategy compared to the constitutive strategy. With the longer cycle time, the optimal facultative strategy resulted in a lower value for *u*\*_*min*_, while *u*\*_*max*_ always remained at 1.

Comparing over all strategies, we found that perfect matching of HIF-α expression to fluctuating levels of oxygenation always produces the highest cell payoff (Fig. 2C). However, for facultative and constitutive strategies that work on non-instantaneous time scales, we found that the superiority of the facultative over the constitutive strategy was low for short cycles and high for longer cycles.

### HIF-α expression under stochastic fluctuations

For stochastic oxygen fluctuations, we convert the cycle times to rates of switching and use these probabilities to create a timeline of stochastic fluctuations comparable to the fractions of time spent in each environment. Specifically, we let P_N→H_ = 1/T_N_ be the probability of switching from normoxia to hypoxia, and P_H→N_=1/T_H_ be the probability of switching from hypoxia to normoxia, and we evaluated the facultative strategy in stochastic environments where P_N→H_ = P_H→N_.

An example simulation with a high probability of switching (left) and a low probability of switching (right) is shown in Fig. 3A. When there is a high probability of switching oxygenation states, P_N→H_ = P_H→N_ = 0.1 min^−1^, the optimal facultative strategy occurs at *u*\*_*min*_= 0.504 and *u*\*_*max*_ = 1. Decreasing the probability of switching to 0.0083 min^−1^ leads to a decrease in *u*\*_*min*_ to 0.216, while *u*\*_*max*_ remains unchanged at 1. For comparison, when P_N→H_ = P_H→N_, assuming *q* = 0.5, the optimal constitutive strategy is u*=0.634. All results for stochastic switching are reported as the mean values of *u*\*_*min*_ and *u*\*_*max*_ for 10 simulations insuring a small standard error (<0.01) for *u*\*_*min*_ and *u*\*_*max*_, respectively (Suppl. Fig. 1).

**Figure 3.**
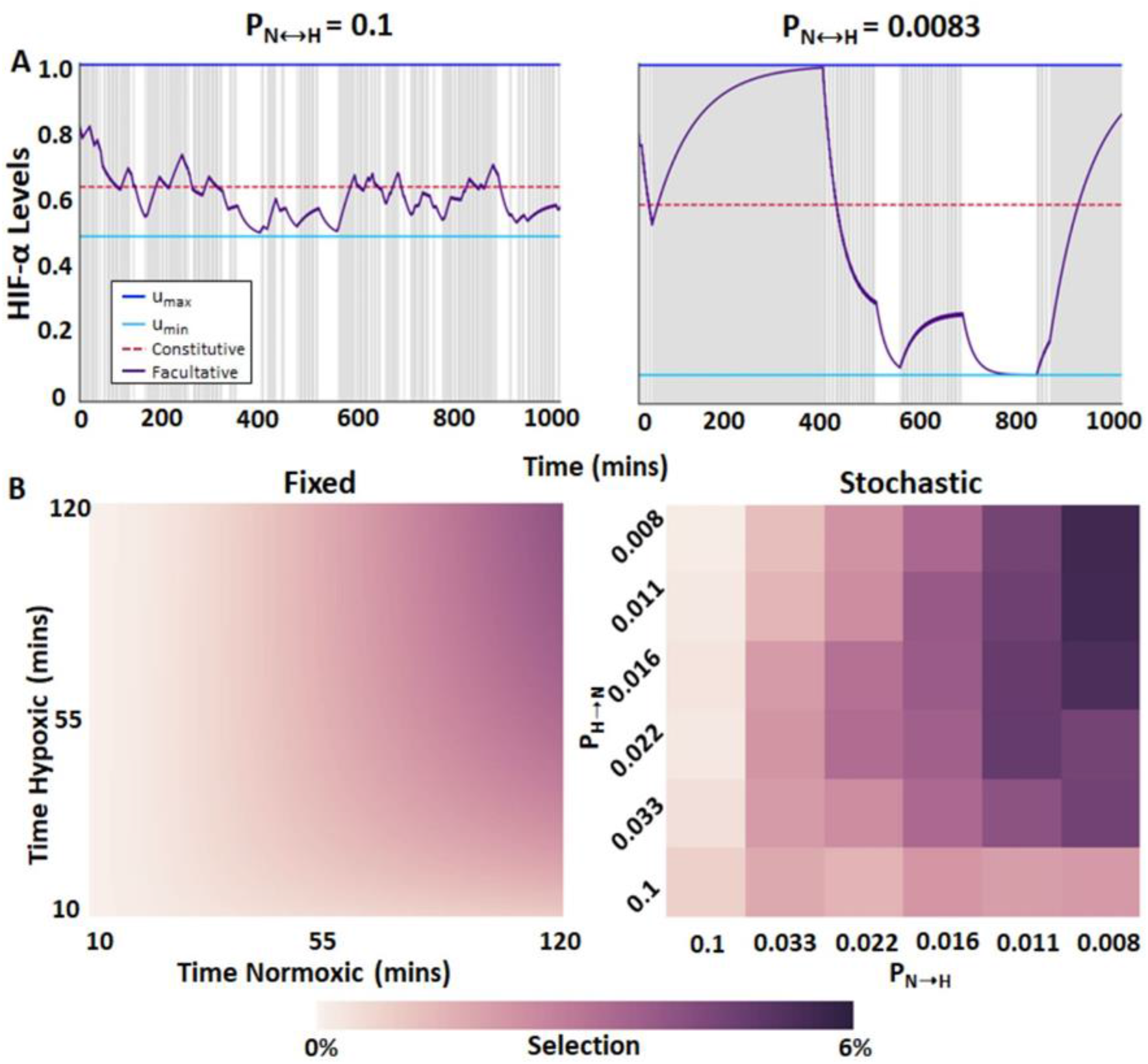
HIF-α regulation under stochastic fluctuations in oxygenation. A) Optimal facultative HIF-α expression in stochastically hypoxic environment; r = 0.00048, α_0_=0.01155, α_1_=0.0462, m = 0.00083, b = 4, c = 0.0001328, and k=1. Left graph illustrates HIF-α stabilization when the probability of fluctuations in oxygenation states is high (P=0.1) and low (P=0.0083). B) Selection for a facultative strategy over a constitutive strategy for fixed interval and stochastic environments. Because selection is optimized numerically in fixed interval environments, selection gradient is continuous. Selection in stochastic environments is presented as the average of 10 simulation results.

### Quantification of stabilization/de-stabilization times in vitro

In the simulations above, we needed estimates for the rates of upregulation and downregulation of HIF-α. We used average rates based on values in the literature from both normal and cancer cell lines. Yet, such rates will vary with cell line and the values from the literature were not collected with our model in mind. To compare to previous values and to our model, we empirically measured HIF-α upregulation and downregulation rates in two different ovarian cancer cell lines, TOV112D and A2780s. The kinetics of HIF-α stabilization were measured under 0.2% hypoxia *in vitro*. For TOV112D, HIF-α upregulation and stabilization required at least one hour and was maximal at 4 hours (Fig. 4A, left panel). For A2780s cells, stabilization required 4 hours (Fig. 4A, right panel). We then measured the length of time required for these cancer cell lines to return to normal HIF-α expression after being exposed to hypoxic conditions for 72 hours. After restoring normoxia, TOV112D returned to normal HIF-α expression in about one minute (Fig. 4B left), while A2780 cells returned to normal HIF-α expression in about five minutes (Fig. 4B right). These are significantly more rapid than prior reports (33-35).

**Figure 4.**
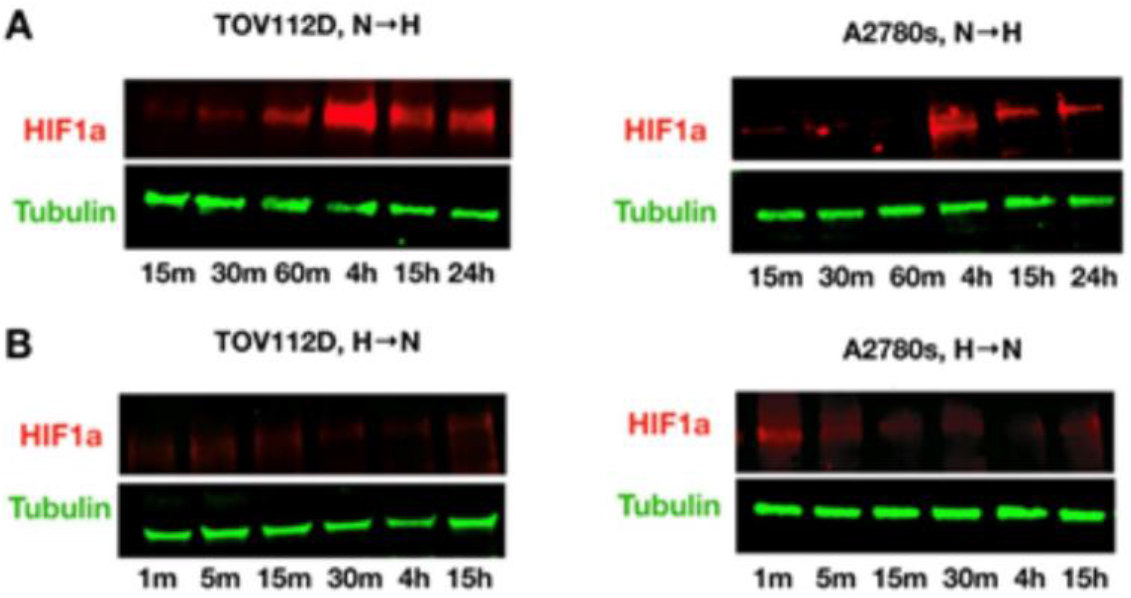
Quantification of HIF-1α stabilization/de-stabilization times in vitro. Using two ovarian cancer cell lines, HIF-1α expression is found in whole lysate by Western blot analysis. Tubulin is used as control of loading the same amount of proteins. A) To measure the stabilization time, cancer cells were cultured in separate dishes, incubated in hypoxia chambers, and collected at several time points. B) To measure the destabilization time, cancer cells were grown for 72h under hypoxia, and collected at several time points after switching to normoxic conditions. See Methods for more details.

We then estimated the upregulation (*α*_0_) and downregulation (*α*_1_) rates of HIF-α. Using Eqs. (S1) and (S2), and assuming that *u*_*max*_=1, *u*_*min*_=0, we estimated the rate of upregulation of HIF-*a* by finding values of *α*_0_ that would allow HIF-α to increase to 90% of its maximum stabilization values (in arbitrary units) in 60 minutes and 240 minutes for TOV112D and A2780 cells, respectively. Similarly, we estimated the rate of HIF-α downregulation by finding values of *α*_1_ that allow HIF-*a* to decline to 10% of its maximum stabilization values within 1 minute and 5 minutes for TOV112D and A2780 cells, respectively. These yielded estimates for *α*_0_∼0.038 min^−1^ for TOV112D and *α*_0_∼0.01 min^−1^ for A2780 and estimates of *α*_1_∼2.3 min^−1^ for TOV112D and *α*_1_∼0.46 min^−1^ for A2780. We incorporated these experimentally derived rates into the model to compare the facultative and constitutive strategies in fixed interval environments. As with our previous results, we find greater selection for a facultative strategy in environments that remain in their current state of oxygenation or lack thereof for longer time periods. The superiority of the facultative strategy over the constitutive depends heavily on up- and down-regulation rates. Decreasing the rate at which HIF-α accumulates reduces the advantage of being facultative over constitutive even in environments with longer cycles. In general, the model suggests that TOV112D cells should exhibit a facultative strategy and A2780 cells a constitutive (see Suppl. Fig. 2).

### Oxygen fluctuations in vivo

Intra-tumoral fluctuations in oxygenation are key for empirically evaluating whether a constitutive strategy may be favored over a facultative one. Furthermore, fluctuations within a tumor may vary spatially. Thus, different regions of a tumor may select for different HIF-α levels and strategies. To gain empirical insights, we measured spatio-temporal variation in oxygen delivery in different regions of a mouse pancreatic adenocarcinoma tumor. We used *in vivo* MR quantification of dynamic T2* changes. Significant heterogeneity was observed within the tumor both in the mean (Fig. 5A) and the temporal variance (Fig. 5B) of T2* values, indicating areas of variable blood flow and oxygenation. The temporal profiles of fluctuations were distinct in different areas, and consistent with different blood vessels feeding these regions. Interestingly, while random, short fluctuations were observed in some areas (Fig. 5C), some displayed changes at much longer time scales (Fig. 5D) in both directions of normoxia to hypoxia and hypoxia to normoxia. We might expect a constitutive strategy to be favored in the former regions and facultative in the latter.

**Figure 5.**
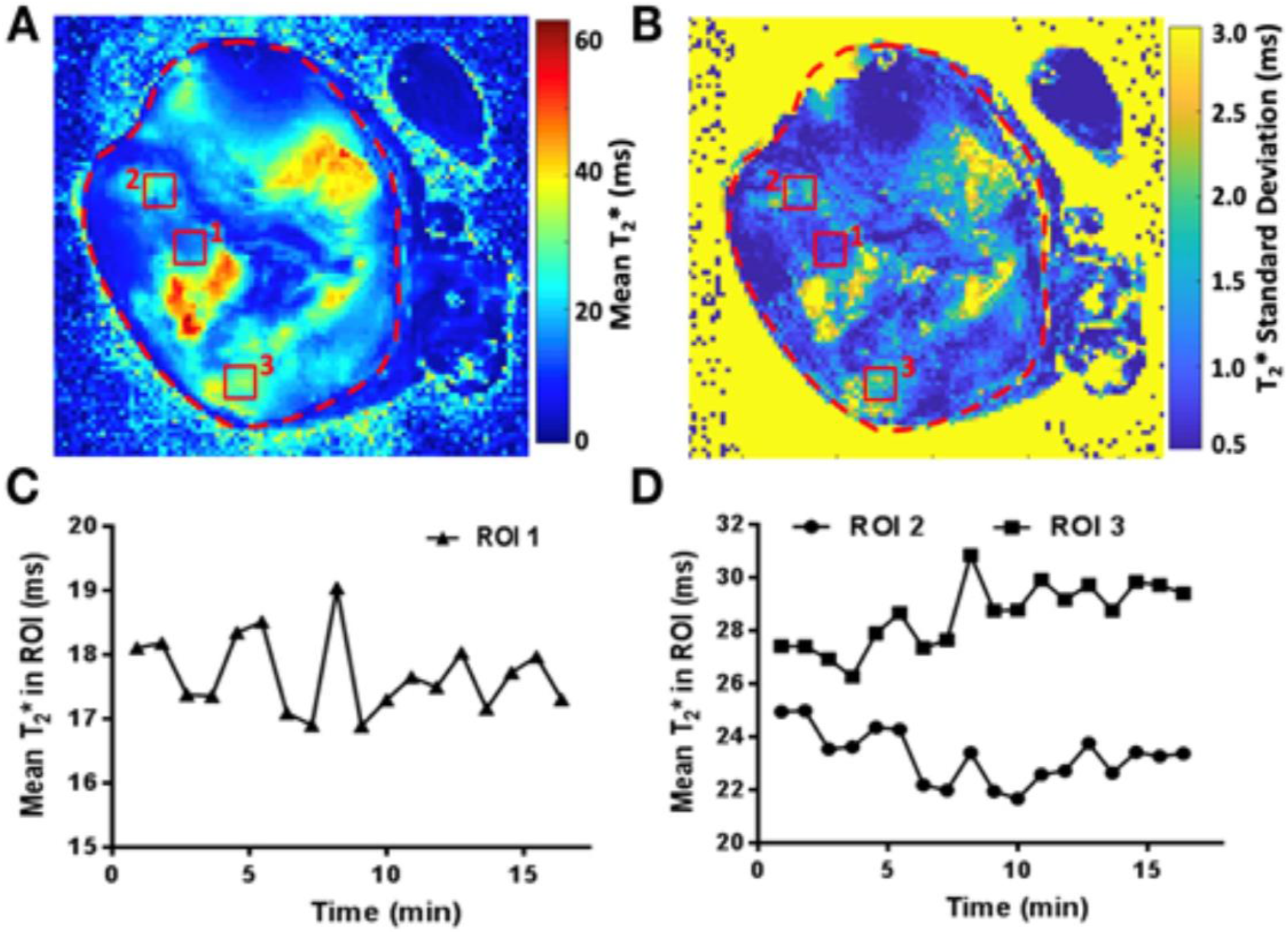
Oxygen fluctuations in vivo. A) Mean T2* value and B) standard deviation of the T2* changes in time are shown for a representative slice. Small regions of interest, marked with red rectangles, were drawn to illustrate distinct temporal T2* kinetics in tumor, plotted in C (ROI 1) and D (ROI 2s and 3).

## Discussion

Facultative regulation of HIF-α in response to fluctuating levels of oxygenation is ancestral and highly conserved across phyla^36-38^. For this reason, the evolution of constitutive HIF-α regulation has drawn wide interest across many biological disciplines, including cancer. Herein, we developed a theoretical model to explore the conditions under which constitutive versus facultative HIF-α orchestration of the cellular response to temporal changes in oxygen supply will optimize a cell’s fitness, as measured by the net growth rate, or payoff. Our modeling indicates, unsurprisingly, that the perfect matching strategy for HIF-α regulation in response to fluctuating oxygenation levels always delivers a greater payoff than either the facultative or the constitutive strategies. However, cellular transcriptional and translational machinery has significant inertia and is unable to instantaneously respond to fluctuations in oxygenation and thus unable to provide a perfect match. Thus, it must respond either facultatively or constitutively. Under facultative regulation, the upper and lower bounds for rates of HIF accumulation are critical for calibrating the rate at which the cancer cells stabilize their oxygen environment. Our model predicts that the upper bound should be set very high to rapidly respond to hypoxia. Conversely the lower bound insures a non-zero level of stabilization. This increased baseline level of stabilization for the facultative strategy is only slightly below what would be expected with constitutive regulation of HIF-α. Importantly, this demonstrates the larger penalty of not upregulating HIF-α quickly enough when the environment becomes hypoxic than the cost incurred of needlessly stabilizing HIF-α during normoxia. Thus, cells constitutively expressing HIF-α may be at a selective advantage under some conditions, with little cost.

Our model indicates that facultative expression of HIF-α always promotes a greater payoff than constitutive expression. Importantly, however, the difference in the payoffs between facultative and constitutive HIF-α expression depends on the nature of the fluctuations between normoxia and hypoxia. With short cycling times, the difference between facultative and constitutive HIF-α stabilization is small and perhaps negligible, so facultative expression may be indistinguishable from constitutive regulation. In contrast, long cycling times or stochastic fluctuations favor facultative HIF-α regulation.

Our modeling results match expectations from nature. For instance, plants rely on inducible (facultative) and/or constitutive defenses against herbivores or pathogens. Optimal defense theory (ODT)^4,32^ predicts deployment strategies for these plant defenses. ODT states that: (1) defenses should be preferentially invested in those tissues that most affect individual fitness, and (2) the reliance on an inducible defense should depend on the probability or predictability of attack (Fig. 6). When the probability of attack is low, there should be greater reliance on inducible versus constitutive defense, and vice-versa when the probability of attack is high. Indeed, as the probability of attack nears 100%, ODT states that fitness is maximized when defenses are constitutive.

**Figure 6.**
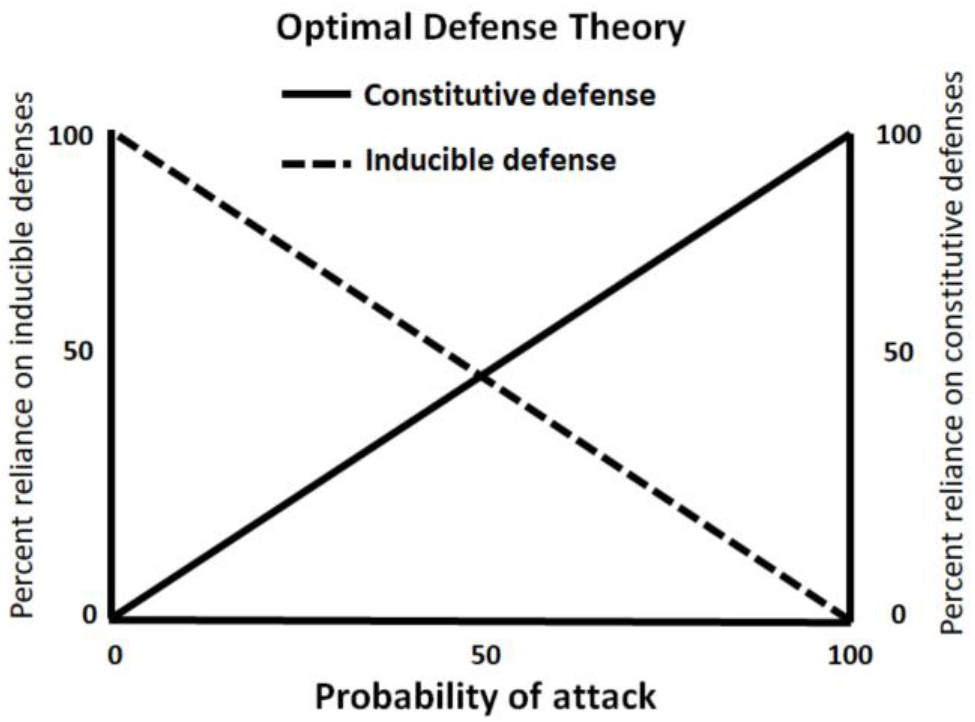
Optimal defense theory. Optimal defense theory predicts that tissues with a low probability of being attacked should rely primarily on inducible defenses, whereas those with a high probability of attack should rely primarily on constitutive defenses.

**Figure 7.**
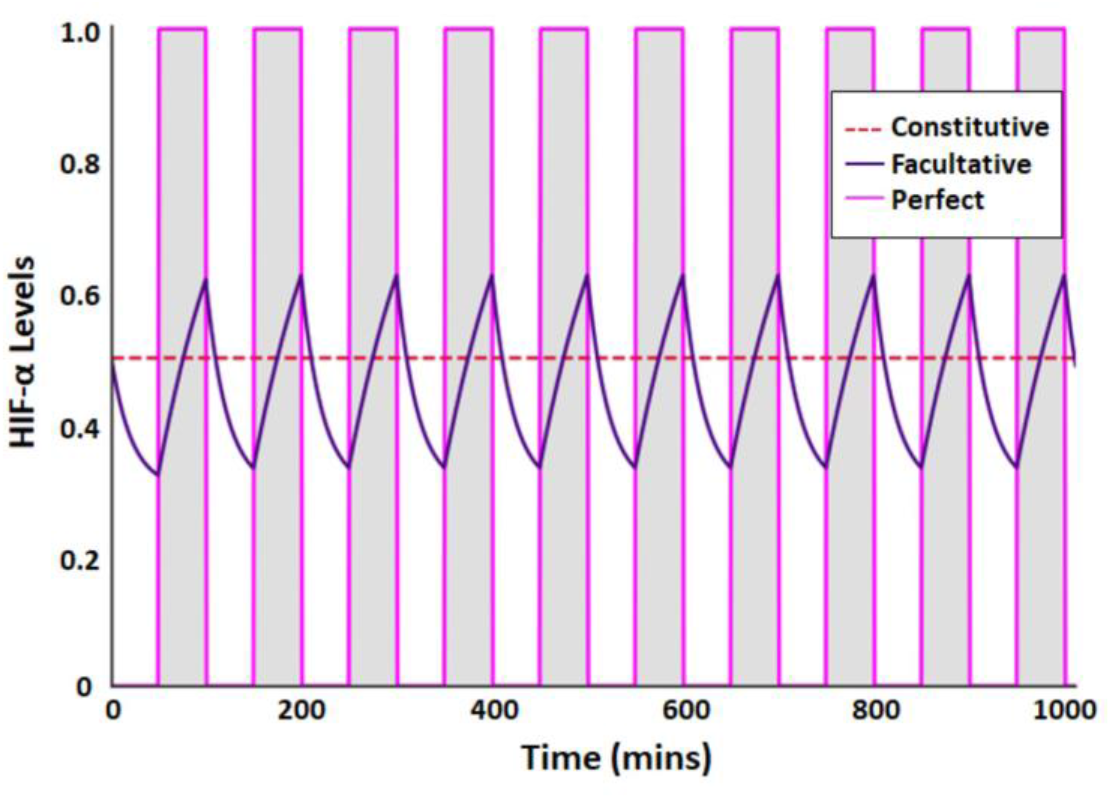
Comparison of 3 HIF-α response strategies to changes in oxygen supply within the tumor (cell) microenvironment. The perfect strategy is instantaneous and used as an upper bound for comparison. Constitutive is constant over time, and facultative has a rate-limited response. The unshaded areas represent periods of full oxygenation while the shaded grey areas represent periods of hypoxia.

Applying this reasoning to HIF-α expression under cyclic hypoxia, fixed periodicities with rapid fluctuations of oxygenation equate to a 100% certainty that a cell will experience hypoxia^39^. Hence, ODT predicts that cells should regulate HIF-α expression constitutively. In our simulations, when oxygenation states switch rapidly at fixed intervals, we found that payoffs are virtually identical between facultative and constitutive HIF-α regulation (Fig 2C). When fluctuations are fixed, and with short periods of normoxia interspersed between long periods of hypoxia, the selective advantage of facultative HIF-α regulation is only slightly greater than with constitutive HIF-α regulation (Fig. 3B). Under both scenarios, any mutation that results in constitutive HIF-α regulation (e.g. mutations in VHL ubiquitin ligase) will be selectively permissive, i.e. it will not be strongly selected against. We speculate that the loss of VHL observed in renal cell cancer implies a constitutive response strategy due to rapid fluctuations in oxygenation early in its development. Cells with a mutation that results in constitutive HIF-α regulation will thus be able to coexist with cells with facultative HIF-α regulation. This mirrors the case in many tumors, in which some cells express the wild-type (normal) HIF-α phenotype (facultative HIF regulation), and some cells express the Warburg phenotype (constitutive HIF-α regulation). Our modeling also suggests that cells evolving under rapid switches in oxygenation may retain facultative HIF-α regulation, but they may set their upper and lower bounds of HIF-α in a way that is effectively pseudohypoxic (Fig. 2B). Under these conditions, the distinction between facultative and constitutive HIF-α regulation becomes moot.

With stochastic fluctuations in oxygenation states, in contrast, the probability of hypoxia is less certain or predictable. Then, as predicted with ODT, facultative HIF-α regulation should be favored. In our simulations, stochastic changes in oxygenation always resulted in greater selection coefficients for facultative relative to constitutive HIF-α stabilization (Fig 3B). Under this scenario, cells with mutations that produce constitutive HIF-α regulation will be less fit than cells with facultative HIF-α regulation, and these cells would be eliminated or reduced to a minor population through competition.

The imaging results presented provide *in vivo* insight into the dynamics of tumor oxygen environment through indirect measurements of blood oxygen level changes in the local vascular network. The apparent fluctuations observed highlight the importance of understanding the cellular mechanisms of adaptation to dynamic conditions. In particular, the measured spatial heterogeneity of the temporal profiles suggests likely coexistence of the different HIF-α regulation strategies within one tumor. With intra-tumoral variation in oxygenation regimes, we expect not only to see variation in the level of HIF-α expression but also the coexistence of cancer cells exhibiting different HIF-α strategies. In the future, spatial relationships can be incorporated into our model to elucidate the nature of these interactions.

Our review of the literature on the dynamics of up- and down-regulation rates, and confirmed by our own experiments indicate that HIF-α response dynamics vary considerably between experimental cell lines (our results above)^33-35^. These differences may represent genetic, epigenetic, or phenotypically plastic differences among tissue types within organisms^40,41^ or evolved differences between species^41,42^. Such heterogeneity may reflect tissue-specific fluctuations in oxygenation, or fluctuations in oxygenation specific to the environments inhabited by different species. This means that cancers initiating from different cell types within different tissues may start with quite varied rates for upregulating and downregulating HIF-α; and these upregulation and downregulation strategies may vary with cancer cell evolution and progression. These differences may later influence the emergence of pseudohypoxia via constitutive HIF-α expression or epigenetic stabilization of downstream products of HIF-α such as CAIX. Of note, though, is the near universality of pseudohypoxia (Warburg Effect) in malignant cancers, indicating that this provides a fitness advantage regardless of the trajectory used to acquire this phenotype^27^.

Previous mathematical models of hypoxia and HIF-α regulation fall into four general categories^43^: (1) understanding the switch-like behavior of the HIF-α response to fluctuating O_2_^44^; (2) analysis of the role of molecular elements of the microenvironment during the HIF-α response^45^; (3) elaborating how asparaginyl hydroxylase factor inhibiting HIF-1 (FIH) affects the HIF-α response^46-48^); and (4) capturing the temporal dynamics of the HIF-α response^46,48^. All of these modeling studies have helped elucidate the core elements that shape the activity and the dynamics of the HIF-α response to cycling hypoxia. Our model, in contrast, addresses the conditions that select for the evolution of constitutive regulation of HIF-α from the ancestral facultative regulation. Our model examines the consequences of different HIF-α regulation strategies on cell fitness within the complex and dynamic tumor ecosystem. We find that constitutive HIF-α regulation is favored when the probability of hypoxia is high – a finding that is consistent with ecological models of defenses to biotic threats like predation and herbivory.

The current study is the first of its kind to apply ecological defense theory to the expression of stress responses (e.g. HIF) in cancer cells. Hence, it is not without its limitations. First, the endpoint for our simulations to model fitness was simply net growth rate, as represented by our payoff function. There are many other components to fitness that were not considered in this study. For example, the expression of some pseudohypoxic gene products, e.g. CAIX or VEGF, may confer upon cells additional selection advantages, such as an increased ability to invade and colonize adjacent tissues^49^, thus increasing the fitness of the pseudohypoxic phenotype. A second limitation is that the study investigated only the kinetics of stabilizing HIF-1α. There are at least two other HIF-α proteins, each with different activation kinetics and portfolios of client proteins. Moreover, the kinetics of the transcriptional and translational machinery induced by HIF-α are not known with certainty, and presumably do not respond instantaneously. Knowledge generated by investigating these limitations will improve subsequent models.

## Methods

### Mathematical Model and Major Assumptions

We modeled three different strategies by which a cell can respond to the presence or withdrawal of oxygen in an environment: perfect, constitutive, and facultative (Fig. 7). If a cell responds perfectly to an environment, the fluctuating oxygen profile would be matched by the cell instantaneously responding with the appropriate HIF-α expression. With constitutive regulation, HIF-α is constantly maintained at an above baseline level regardless of the O_2_ levels. With facultative regulation, HIF-α levels change in response to environmental fluctuations in oxygen at fixed rates of accumulation (slow) and degradation (fast). It is important to note that we use “HIF-α” to represent the constellation of cellular responses to hypoxia. While the transcription factor HIF-1α is undoubtedly central to this, we do not wish to imply that its levels are solely responsible for a cell’s fitness under different conditions of oxygenation.

### Environment creation

Let *Y* ∈ [0, 1], describe the level of oxygenation in the cell’s tumor microenvironment. In all scenarios, we assume that the environmental switch between fully oxygenated (*Y*=1) or deoxygenated (*Y*=0) is effectively instantaneous whereas the accumulation and degradation of HIF-α is based on intrinsic rates. Let *u*(*t*) be the HIF-α response of a cell at time *t* whether it has a perfect, constitutive, or facultative response, where *u* ∈ [0, 1]. For the fixed interval environment, normoxic (Y=1) and hypoxic (Y= 0) periods switch back and forth at fixed time intervals. A perfect, constitutive, and facultative response to these changing oxygen profiles will exhibit different HIF-α expression levels. (Fig. 7).

### The perfect strategy

With the perfect strategy, the switching response to the environment is instantaneous. Therefore, *u*=0 during periods of normoxia and *u*=1 during periods of hypoxia. We can thus simply take the sum of the payoffs spent in each environment using Eq. (1), and expected payoff becomes:

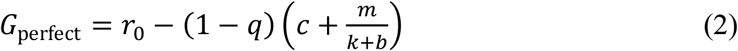

### The constitutive strategy

With the constitutive strategy, HIF-α levels remain constant and do not change in response to the temporal fluctuations in oxygen. When *u* is fixed, the expected payoff for the constitutive HIF-α strategy can be separated into the payoff during normoxia, *G*_N_(*u*) = *r* - *cu*, and the payoff during hypoxia, *G*_H_(*u*) = *r* - *cu*-*m*(1-*q*)/(*k*+*bu*), so that the total payoff is:

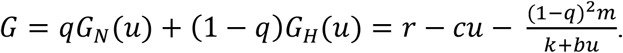

The optimal value for HIF-α expression depends only on the fraction of time spent in each state, *q*. Therefore, the optimal constitutive strategy value, *u*\*, can be calculated analytically by maximizing expected *G* with respect to *u*. We take the first order necessary condition for this optimum, 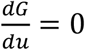, and solve for u*, resulting in:

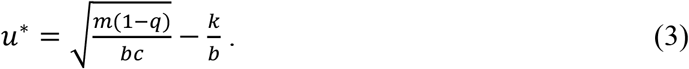

The constitutive level of HIF-α production declines with its proliferation cost, *c*, the cell’s intrinsic tolerance to hypoxia in the absence of HIF-α stabilization, *k*, and the fraction of time spent normoxic, *q*. HIF-α production increases with cell mortality when conditions are hypoxic, *m*. The relationship between optimal HIF-α production and the benefit of HIF-α expression *u* in reducing mortality when conditions are hypoxic, *b*, is hump shaped. If HIF-α expression is ineffective (small *b*) then there is no point, and if HIF-α expression is extremely effective (large *b*) then little is needed. The *u*\* can then be substituted into the payoff *G* to determine the maximal payoff available to the constitutive strategy given the micro-environmental and fitness parameters:

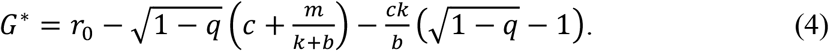

### The facultative strategy

Under the facultative strategy, when the environment is depleted of oxygen, we assume that the cell accumulates HIF-α at a finite rate *α*_*0*_, and when the environment is fully oxygenated the cell degrades HIF-α at a finite rate *α*_*1*_ (1). We allow a baseline production of HIF-α (*u*_*min*_) even under normal oxygen conditions and assume that the cell targets a maximum accumulation of HIF-α (*u*_*max*_) in an oxygen-depleted environment (2). We assume that the changes in expression occur at a rate proportional to the difference between some desired level and the current level, such that:

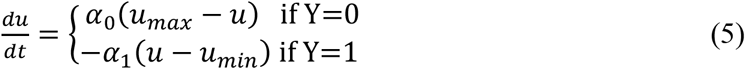

For the facultative strategy, the dynamic *u*(*t*) requires that the expected payoff be calculated as the cumulative payoff over the normoxic and hypoxic periods, *T*_*N*_ and *T*_*H*_, respectively. Because the HIF-α fluctuations become periodic, the total cumulative payoffs can be averaged over a cycle separated into the time spent in normoxia and the time spent in hypoxia:

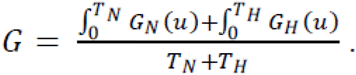

The payoff can be analytically solved to:

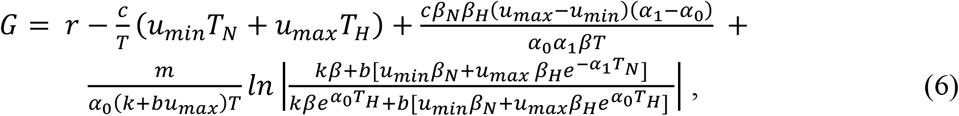

where 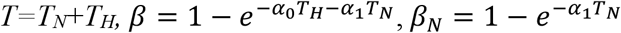, and 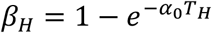. The derivation is provided in the supplemental material.

The payoff for the facultative HIF-α strategy in a stochastic environment is calculated similarly. But, because the HIF-α dynamics cannot settle into a dynamic equilibrium, the payoff is calculated over discretized time intervals and then taken as an average over the entire simulation.

### Finding the optimal u*_min_ and u*_max_ for the facultative strategy

Constrained optimization by linear approximation (COBYLA) was used to determine values of *u*\*_*min*_ and *u*\*_*max*_ that maximize a cell’s payoff. For stochastic environments, a search space was created by linearly separating 250 values between 0 and 1. These values were used to create combinations of *u*_*min*_ and *u*_*max*_ for which 0 ≤ *u*_*min*_ < *u*_*max*_ ≤ 1. After *u*(*t*) was computed for each combination of *u*_*min*_ and *u*_*max*_, the payoff, *G*(*u*(*t*)), was calculated as an average of all payoff values at each time point. The combination of *u*_*min*_ and *u*_*max*_ that produced the maximum averaged payoff was considered optimal. For a given P_N→H_ = P_H→N_, we ran 10 replicate runs for 2000 time units. With this number of time units, the estimated values of *u*\*_*min*_ and *u*\*_*max*_ were very similar across replicate runs (standard error of the mean < 0.01).

### Selection coefficient for facultative expression

We define the selection coefficient (SC) as the fitness advantage for using a facultative rather than a constitutive strategy. We calculated the SC as the difference between the payoff for facultative expression, *G*_*F*_, and the payoff for constitutive expression, *G*_*C*_, normalized by the payoff for constitutive expression, *SC*=(*G*_*F*_-*G*_*C*_)/*G*_*C*_.

### Cell lines and Culture Conditions

A2780s and TOV112D ovarian cancer cells were obtained through American Type Culture Collection (ATCC). Cells were grown in RPMI supplemented with 10% fetal bovine serum (FBS). For both normoxic and hypoxic treatment environments, all cells were grown in 5% CO_2_ and at 37°C in a humidified atmosphere.

### Stabilization/degradation of HIF

A Biospherix X-Vivo Hypoxia Chamber was used to incubate cells under hypoxic conditions. For hypoxic conditions, cells were incubated at 0.2% O_2_:94.8% N_2_:5% CO_2_. Reoxygenation was performed by transferring flasks or plates containing cells from the hypoxic chamber to an incubator under atmospheric conditions at 5% CO_2_. For time points shorter than 4 hours, media pre-equilibrated under hypoxic conditions was used. Then, the hypoxic media was added to the cells inside a hypoxic (0.2% O_2_) working chamber within the Biospherix complex.

### Western blot analyses

Western blots were performed on A2780 and TOV112D ovarian cancer cells to validate the expression of HIF at the protein level at different time points. Cells grown in hypoxic chambers were frozen at the time points mentioned in the results section and harvested all together by lysing in Radioimmunoprecipitation assay buffer (RIPA buffer) containing 1× protease inhibitor cocktail (Sigma-Aldrich). For each sample, a 30 μg aliquot was loaded onto pre-cast polyacrylamide-SDS gels from BioRAd that were then transferred onto nitrocellulose. Membranes were incubated with primary antibodies against HIF-1α (#610958, BD Biosciences), Tubulin (#2144, CST) or β-Actin (A5441, Sigma, 1:4000) overnight at 4° C, followed by fluorescent-conjugated secondary antibodies (IRDye^®^ 800CW Goat anti-MouseIgG and IRDye^®^ 8680CW Goat anti-rabbit IgG). An Odyssey chemiluminescence-fluorescence system was used for protein detection. Proteins detected ran at the expected molecular weights, as verified using molecular weight standard markers.

### MRI tumor imaging of hypoxia

*In vivo* measurements of oxygenation fluctuations were obtained by Intrinsic Susceptibility Magnetic Resonance Imaging (IS-MRI)^50^. Panc02 mouse pancreatic adenocarcinoma cells were implanted subcutaneously into a C57BL/6 mouse. When the tumor reached a volume of ∼1500mm^3^, as measured by calipers, the animal underwent MR imaging with a 7T/30cm Bruker Biospec® imaging spectrometer as follows. Mice were anaesthetized using 3% isoflurane and subsequently maintained with 1.5-2% isoflurane mixed with 100% oxygen. Anatomical images were acquired using T2-weighted coronal and axial scans to identify the middle of the tumor and facilitate outlining the tumor. To capture the spatial and temporal dynamics of oxygenation, quantitative T2* mapping was then performed continuously every 55s for 18 series (Multi-Gradient Echo, TR=270ms, 10xTE=2.5-47.5ms, flip angle 40 degrees, 5 slices, 1mm/1mm slice thickness/gap, 35mm FOV, 128×128 points). A monoexponential function was fitted for each voxel at each time-point (MATLAB 2018b, Mathworks) to reconstruct the local T2* magnetic resonance time, which is modulated by changes in deoxyhaemoglobin levels in the blood, hence reflecting the fluctuations in blood oxygen level.

## Acknowledgements

This work was supported by NIH/NCI PSOC U54CA193489, “Cancer as a Complex Adaptive System”, NIH/NCI P30CA076292, “Moffitt Cancer Center Support Grant”, and NIH/NCI U54 Supplement, “The tumor-host evolutionary arms race”.

## Author contributions

R.J.G., J.S.B., and C.J.W. conceived of and designed the research. M.P., J.A.G. and J.S.B. performed the mathematical modeling. M.R.T., M.D. and P.B. conducted the laboratory work. M.P., J.A.G, and C.J.W. wrote the manuscript, and M.P, J.A.G., J.S.B., M.R.T., M.D., R.J.G. and C.J.W. revised the manuscript.

