## Supplemental Material for "Cycling hypoxia selects for constitutive HIF stabilization"

**Cycling hypoxia selects for constitutive HIF stabilization --Supplemental Material**

**Derivation of Eq. 5.**

Solving Eq. (5) for $u$ when the environment is normoxic (*Y*=1) we obtain

$u_{N}(t)=u_{min}+e^{-\alpha_{1}t}(u_{N}\left( 0 \right)-u_{min})$ , (S1)

and when the environment is hypoxic (Y=0) we obtain

$u_{H}(t)=u_{max}+e^{-\alpha_{0}t}(u_{H}\left( 0 \right)-u_{max})$ . (S2)

These two equations represent the dynamics of HIF-α levels. If the fluctuations are periodic, then the HIF-α level at the end of a period of normoxia would equal the initial HIF-α level when the environment becomes hypoxic, $u_{N}\left( T_{N} \right)=u_{H}\left( 0 \right)$, and vice versa, $u_{H}\left( T_{H} \right)=u_{N}\left( 0 \right)$. These equalities can be simplified to

${u_{N}\left( 0 \right)=u}_{max}+e^{-\alpha_{0}T_{H}}\left( u_{H}\left( 0 \right)-u_{max} \right)$	(S3)

$u_{H}\left( 0 \right)=u_{min}+e^{-\alpha_{1}T_{N}}\left( u_{N}\left( 0 \right)-u_{min} \right)$ . (S4)

We can substitute Eq. (S4) into Eq. (S3) and Eq. (S3) into Eq. (S4) to solve for $u_{N}\left( 0 \right)$ and $u_{H}\left( 0 \right)$, respectively, yielding

$u_{N}\left( 0 \right)=\frac{u_{max}(1-e^{-\alpha_{0}T_{H}})+u_{min}e^{-\alpha_{0}T_{H}}(1-e^{-\alpha_{1}T_{N}})}{1-e^{-\alpha_{0}T_{H}-\alpha_{1}T_{N}}}$	(S5)

$u_{H}\left( 0 \right)= \frac{u_{min}(1-e^{-\alpha_{1}T_{N}})+u_{max}e^{-\alpha_{1}T_{N}}(1-e^{-\alpha_{0}T_{H}})}{1-e^{-\alpha_{0}T_{H}-\alpha_{1}T_{N}}}$ . (S6)

We can then solve for the payoff during normoxic time periods using Eq. (1) from the main text, which for normoxia simplifies to $G_{N}(t)=r-cu_{N}(t)$. Substituting in Eq. (S1) gives

$G_{N}(t)=r-c[u_{min}+e^{-\alpha_{1}t}(u_{N}\left( 0 \right)-u_{min})]$ . (S7)

Then the total payoff during a normoxic period is found by integrating $G_{N}$ from 0 to *T_N_*

$\int_{0}^{T_{N}} G_{N}=rT_{N}-{cu}_{min}T_{N}+\frac{c\left( 1-e^{-\alpha_{1}T_{N}} \right)}{\alpha_{1}}\left[ u_{min}-u_{N}\left( 0 \right) \right]$ . (S8)

Finally, substituting Eq. (S5) in Eq. (S8) we obtain

$\int_{0}^{T_{N}} G_{N}=rT_{N}-{cu}_{min}T_{N}+\frac{c{\beta_{H}\beta}_{N}{(u}_{min}-u_{\max})}{\alpha_{1}\beta}$ , (S9)

which is simplified by 𝛽$=1-e^{-\alpha_{0}T_{H}-\alpha_{1}T_{N}}$, $\beta_{N}=1-e^{-\alpha_{1}T_{N}}$, and $\beta_{H}=1-e^{-\alpha_{0}T_{H}}$.

The method is the same for the hypoxic time periods, however, the payoff is slightly more complicated by the mortality term: $G_{H}(t)=r-cu_{H}\left( t \right)-\frac{m(1-q)}{k+bu_{H}(t)}$ . Eq. (S2) can be substituted into the payoff equation to obtain

$G_{H}(t)=r-c[u_{max}+e^{-\alpha_{0}t}(u_{H}\left( 0 \right)-u_{max})]- \frac{m}{k+b\left[ u_{max}+e^{-\alpha_{0}t}(u_{H}\left( 0 \right)-u_{max}) \right]}$ . (S10)

Then we find the total payoff during a hypoxic interval by integrating $G_{H}$(t) from 0 to *T_H_*, as

$$\int_{0}^{T_{H}} G_{H}=rT_{H}-cu_{max}T_{H}+\frac{c\left( 1-e^{-\alpha_{0}T_{H}} \right)}{\alpha_{0}}\left[ u_{max}-u_{H}\left( 0 \right) \right]$$

(S11)

$+\frac{m}{\alpha_{0}\left( k+bu_{max} \right)}ln\left| \frac{k+bu_{H}\left( 0 \right)}{e^{\alpha_{0}T_{H}}(k+bu_{max})-b\left[ u_{max}-u_{H}\left( 0 \right) \right]} \right|$ .

And finally, we substitute Eq. (S6) into Eq. (S11) to obtain

$$\int_{0}^{T_{H}} G_{H}=rT_{H}-cu_{max}T_{H}+\frac{c\beta_{H}\beta_{N}\left( u_{max}-u_{min} \right)}{\alpha_{0}\beta}$$

(S12)

$+\frac{m}{\alpha_{0}\left( k+bu_{max} \right)}ln\left| \frac{k\beta+b(u_{min}\beta_{N}+u_{max}{\beta_{H}e}^{-\alpha_{1}T_{N}})}{k\left( e^{\alpha_{0}T_{H}}-e^{-\alpha_{1}T_{N}} \right)+b(u_{min}\beta_{N}+u_{max}\left( e^{\alpha_{0}T_{H}}-1 \right))} \right|$,

again simplifing by 𝛽$=1-e^{-\alpha_{0}T_{H}-\alpha_{1}T_{N}}$, $\beta_{N}=1-e^{-\alpha_{1}T_{N}}$, and $\beta_{H}=1-e^{-\alpha_{0}T_{H}}$.

The payoff over a complete cycle is expressed as the sum of the payoff over normoxia (Eq. (S9)) and the payoff over hypoxia (Eq. (S12)), divided by total time $T= T_{N}+T_{H}$. After some rearrangement, this is simplified to

$G= r-\frac{c}{T}\left( u_{min}T_{N}+u_{max}T_{H} \right)+\frac{c\beta_{N}\beta_{H}\left( u_{max}-u_{min} \right)\left( \alpha_{1}-\alpha_{0} \right)}{{\alpha_{0}\alpha}_{1}\beta T} +$

$\frac{m}{\alpha_{0}\left( k+bu_{max} \right)T}ln\left| \frac{k\beta+b[u_{min}\beta_{N}+u_{max}\beta_{H}e^{-\alpha_{1}T_{N}}]}{k\beta e^{\alpha_{0}T_{H}}+b[u_{min}\beta_{N}+u_{max}{\beta_{H}e}^{\alpha_{0}T_{H}}]} \right|$ . (S13)

**Supplemental Figure 1**

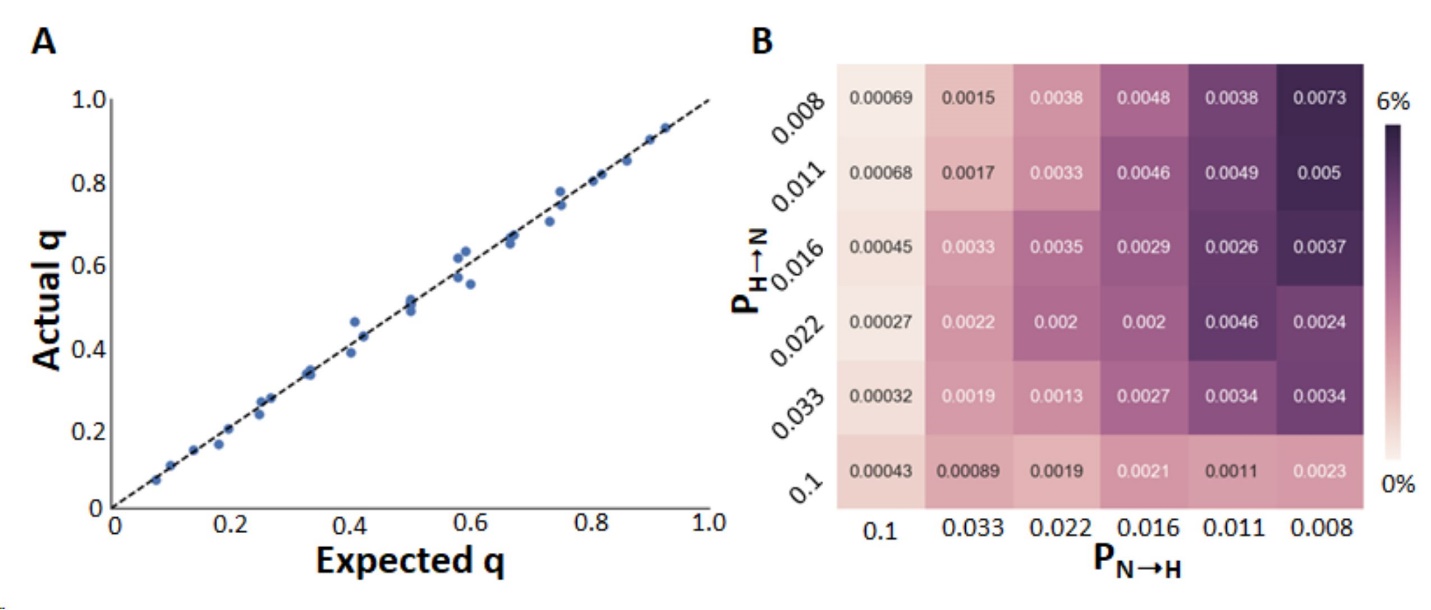

**Supplemental Figure 1.** The average results of 10 stochastic simulations. A) In each simulation, the probability that the environment will remain or switch its state of oxygenation is determined stochastically. We plot the expected probability that the environment will be normoxic (expected q) against the actual q, calculated after the environment is produced. B) Selective advantage of facultative HIF-α regulation in stochastic environments. Each cell is annotated with the standard error.

**Supplemental Figure 2**

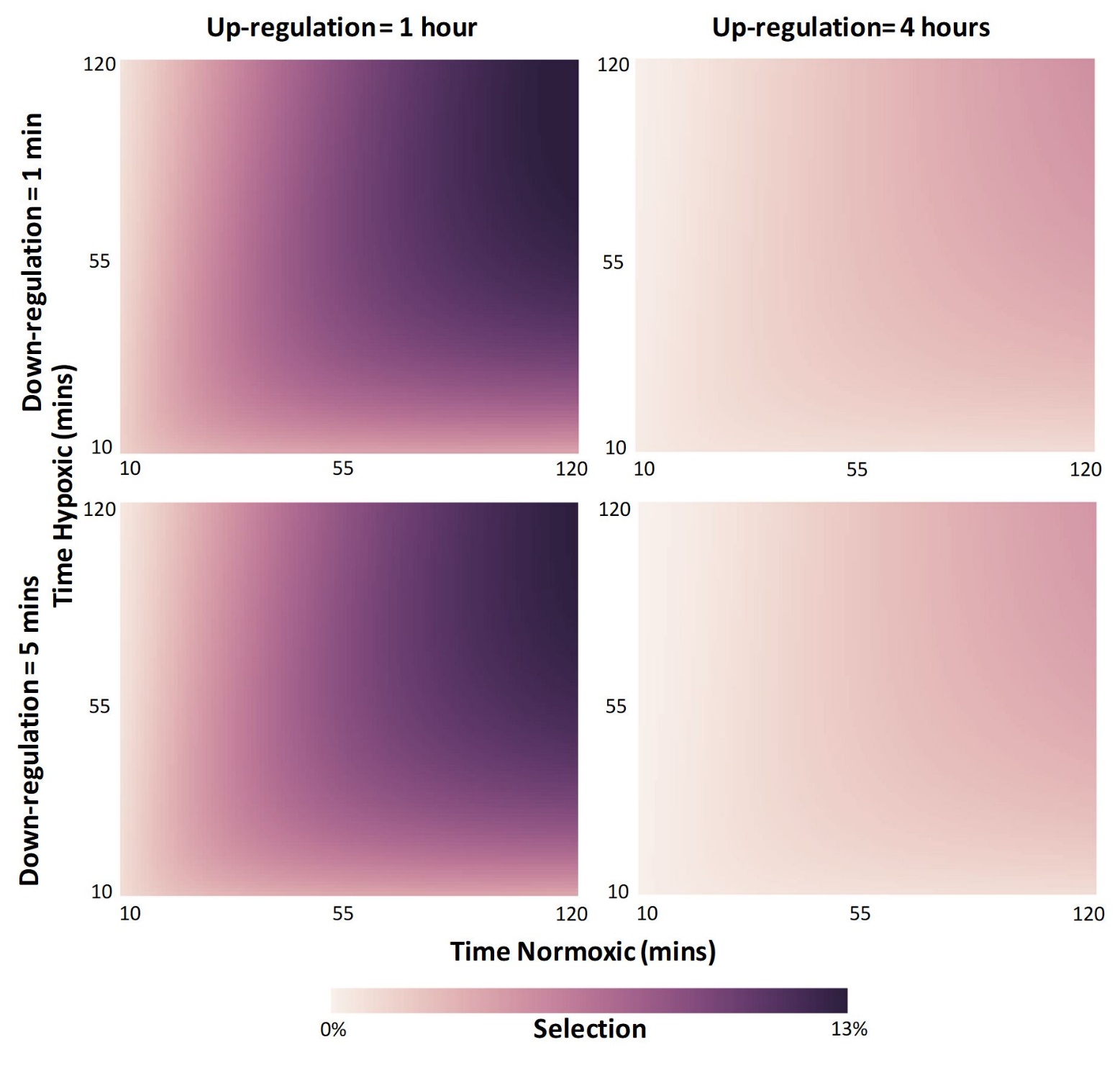

**Supplemental Figure 2.** Selective advantage of facultative versus constitutive HIF-α regulation in environments with fixed intervals of normoxia and hypoxia when the rates of HIF-α accumulation (α_0_) and degradation (α_1_) differ. For each subgraph, values of α_0_ and α_1_ were chosen to produce accumulation and degradation times indicated. For accumulation of 1 hour, α_0_ = 0.038 min^-1^ and for accumulation of 4 hours, α_0_ = 0.01 min^-1^. For degradation of 1 min, α_1_ = 2.3 min^-1^ and for degradation time of 5 min, α_1_ = 0.46 min^-1^.
